# High resolution evolutionary analysis of within-host hepatitis C virus infection

**DOI:** 10.1101/400283

**Authors:** Jayna Raghwani, Chieh-Hsi Wu, Cynthia K. Y. Ho, Menno de Jong, Richard Molenkamp, Janke Schinkel, Oliver G. Pybus, Katrina A. Lythgoe

## Abstract

Despite the breakthroughs in the treatment of HCV infection in recent years, we have a limited understanding of how virus diversity generated within individuals impacts the evolution and spread of HCV variants at the population scale. Addressing this gap will be important for building models for molecular epidemiology, which can identify main sources of disease transmission and evaluate the risks of drug-resistance mutations emerging and disseminating in a population. Here, we have undertaken a high-resolution analysis of HCV within-host evolution from four individuals co-infected with HIV. Specifically, we used long-read, deep-sequenced data of the full-length HCV envelope glycoprotein, longitudinally sampled from acute to chronic HCV infection to investigate the underlying viral evolutionary dynamics. In three individuals we found strong statistical support for population structure maintaining within-host HCV genetic diversity. Furthermore, we found significant variation in rates of molecular evolution among different regions of the HCV envelope region, both within and between individuals. Lastly, we report the first estimate of the within-host population genetic rate of recombination for HCV (0.28 x 10^-7^ recombinations per site per day; interquartile range: 0.13-1.05 x 10^-7^), which is two orders of magnitude lower than that estimated for HIV-1, and four orders of magnitude lower than the nucleotide substitution rate of the HCV envelope gene. Together, these observations indicate that population structure and strong genetic linkage shapes within-host HCV evolutionary dynamics. These results will guide the future investigation of potential HCV drug resistance adaptation during infection, and at the population scale.

## INTRODUCTION

Hepatitis C virus (HCV) is a fast-evolving RNA virus that is estimated to infect 80 million people globally (1). Although the virus is spontaneously cleared in 15-30% of newly infected patients, remaining individuals become chronically infected, potentially leading to severe liver disease, such as cirrhosis and liver cancer. The recent rollout of direct-acting antiviral (DAA) drugs against HCV, which can cure infection in more than 90% of cases and have considerably less toxicity than previous drug treatment regimens (2), has been a major breakthrough. However, treatment success varies among viral strains, host genotype, and disease status (3, 4). In addition, DAAs remain out of reach for many individuals, especially if they are unaware of their infection and/or are living in resource-poor countries with high HCV burdens. Moreover, even though DAAs are highly effective in clearing viral infection, they do not provide long-term protection against reinfection with HCV, which is a likely occurrence in risk groups, such as HIV-positive men who have sex with men, with high rates of exposure to the virus (5).

In order to achieve the WHO target of eliminating HCV by 2030, it will be necessary to focus on developing the most effective strategies for targeted treatment and intervention. Molecular epidemiology has become a powerful tool for understanding how viruses spread through populations, identifying predominant sources of disease transmission, and developing public health interventions (6). This approach has been commonly employed for HIV-1 (e.g. (7)), but comparable studies for HCV are more challenging due to the high viral diversity observed during HCV infection, which makes it difficult to link infected individuals phylogenetically. In particular, there is a growing body of work indicating that within-host HCV diversity is maintained by multiple viral subpopulations with independent replication behaviour, where distinct long-lived lineages are intermittently detected in blood plasma over time (8-12). This observation implies that transmitted viruses are unlikely to be representative of the donor within-host HCV population sampled at a different point in time, and suggests there is potential for divergent viral strains to be transmitted from the same source individual. Consequently, to build more effective molecular epidemiology approaches for HCV, we need to understand what processes shape the within-host viral diversity, and how this impacts the evolution and spread of viral variants at the population scale.

Insights into within-host HCV evolutionary dynamics have typically been gained from short regions of the virus genome (often ∼300bp) and from a small number of viral sequences sampled during infection (8, 10, 13-16). Consequently, it has been difficult to test directly and with statistical confidence if the observed HCV molecular evolutionary patterns are consistent with viral population structure, in contrast to a single viral population that undergoes repeated bottlenecks (i.e. fluctuating in population size over time). Furthermore, although ultra-deep short-read sequence data are now commonly generated in studies of within-host viral evolution (e.g. to detect low-frequency viral variants and estimate transmission bottlenecks) such data are less suitable for reconstructing infection dynamics from sequences due to the large phylogenetic uncertainty associated with short reads. To address these challenges, we undertook an evolutionary analysis of full-length HCV envelope (E1/E2) gene sequences serially sampled from four patients, which were generated previously using Pacbio SMRT sequencing (12). The high resolution provided by long-read, high-throughput, serially-sampled virus sequences is ideal for studying the within-host evolution of chronic viruses, such as HIV and HCV, which are characterised by complex and genetically diverse viral populations. Importantly, by sequencing complete gene regions at greater depth, we have sufficient statistical power to distinguish viral variants that have emerged and persisted over the course of infection, enabling more detailed insights into within-host viral evolutionary dynamics.

We first investigated HCV population dynamics from acute to chronic infection, and asked whether the observed patterns support a structured population with at least two independently replicating viral subpopulations, or a single population that fluctuates in size over time. For three out of four individuals, we found strong evidence for within-host viral population structure. Next, we estimated the evolutionary rates of the HCV envelope glycoprotein during infection, which indicated considerable variation in evolutionary rate among the gene regions within E1E2, and among individuals. Finally, we calculated genetic divergence over the course of infection, and the estimated the within-host recombination rate. Notably, we conclude that HCV recombines considerably less frequently than HIV-1. This is in line with the observation that HCV has fewer recombinant forms circulating at the epidemiological level than HIV-1, even though HCV is more transmissible and mixed infections are very common.

## METHODS

### Sequence data

We analysed full-length HCV envelope (1680 bp) sequences from four individuals that were coinfected with HIV-1 and HCV. The individuals were infected with HCV genotype 4d and were sampled longitudinally from acute to chronic infection for durations between 5 to 12 years, at average intervals of 0.66 to 1.10 years. The sequence data were generated by PacBio SMRT sequencing using stringent multiple passes to infer the circular consensus sequence (CCS) reads. This approach reduced the consensus sequence error rate from 1.6% to 0.4% and resulted in approximately 500 and 740 CCS reads per sample (12). Further information about the sequencing and individual samples can be found in C. K. Ho, et al. (12).

### Evolutionary Dynamics of HCV infection

Time-scaled phylogenies and demographic histories of within-host HCV infections were estimated in BEAST v.1.8 (17) using a relaxed uncorrelated log-normal distributed molecular clock (18), a codon-structured nucleotide substitution model (19), and a Bayesian Skygrid coalescent prior (20). Bayesian phylogenetic inference approaches, such as those implemented in BEAST, are computationally prohibitive for large sequence datasets (>2000 sequences). Therefore, we randomly subsampled 25 reads per time-point per individual, resulting in datasets ranging from 225 to 300 sequences per individual. Two to four independent MCMC chains of 200 million steps were computed for each data set (one data set per individual) to ensure that adequate MCMC mixing and stationarity had been achieved. The resulting phylogenies were plotted with ggtree (21), and trends in within-host HCV population dynamics were inferred with ggplot2 (22) by fitting a loess regression through the N_e_τ values over time (estimated from the Bayesian Skygrid model).

The posterior trees from the above analysis were subsequently used as empirical tree distributions for estimating codon-partitioned substitution rates (CP1+2: first and second codon positions and CP3: third codon position) in BEAST v1.8 (17) which were used as proxies for the nonsynonymous and synonymous substitution rates. The alignment was partitioned into three main subgenomic regions, (i) E1, (ii) E2 excluding the hypervariable region 1 (HVR1), and (iii) HVR1. Two independent MCMC chains of 10 million steps were computed for each individual data set to ensure adequate mixing and stationarity had been achieved.

To investigate whether the pattern of population genetic diversity during HCV infection is better explained by a structured within-host population characterised by two independently replicating populations, or by a single population with temporally varying population size, we compared two coalescent models: an approximation of the structured coalescent (23) and the Bayesian skyline coalescent (24). To avoid any biases that may be produced from unequal sampling through time, datasets were subsampled such that each individual dataset contained an equal number of sequences in total (25). For the structured coalescent analysis, two demes were defined such that one corresponded to an observed population in the blood and the other to an unobserved population, which we posit exists most likely within the liver, but could represent other compartments (9, 11). Crucially, we assume that both subpopulations originate from the liver, since this is the main site of HCV replication, but only one of these subpopulations is detected in the blood at any given sampling event (9, 11). As all HCV sequences were sampled from the blood, we constrained the between sub-population migration rates to be equal. Biologically, this corresponds to viral lineages moving between the liver and blood (i.e. where viruses are shed into the blood from the liver) at equal rates. To compare the two coalescent models, their corresponding marginal likelihoods were calculated using the stepping-stone sampling method (26), with an alpha parameter of 0.3 and 8 steps. The calculations were performed using the MODEL-SELECTION package of BEAST2 (27). Since the marginal likelihood calculations require proper priors (i.e. the prior distribution must integrate to 1), the Bayesian skyline coalescent prior was chosen instead of the Bayesian skygrid coalescent prior for estimating the marginal likelihood of a single population whose size varies through time. We assumed equal prior probabilities on Bayesian skyline and BASTA, as the aim was to determine which models fits the data better for each patient. Therefore, comparison by the marginal likelihood values directly is equivalent to using Bayes factors. Greater support for a structured within-host population is indicated if a higher marginal likelihood is observed with BASTA than for the Bayesian skyline coalescent model, and vice versa.

### Divergence over time

For each patient, the full set of sequences (i.e. not subsampled) was used to estimate the nucleotide divergence of the within-host HCV population over time using a custom Python script (https://github.com/jnarag/HCVPacbioAnalysis.divergence.py). Specifically, divergence at a given time-point was calculated as the average per-site nonsynonymous or synonymous nucleotide difference compared to the founder viral strain. The founder viral strain used was the consensus virus sequence at the first time-point. The per-gene divergence was estimated by partitioning the alignments according to the three subgenomic regions, as described above.

### Per-site amino-acid diversity

The mean per-site amino-acid diversity was estimated for each patient using full set of sequences with a custom-made python script (https://github.com/jnarag/HCVPacbioAnalysis/AAdiversity.py). Only sites that contained no gaps were considered. Mean amino-acid diversity was calculated as the number of amino-acid differences among all pairs of sequences divided by the total number of unique pairs of sequences in each alignment.

### Within-host HCV recombination rate

The average per site per year effective recombination rate across the four individuals was estimated using a custom-made Python script based on the approach outlined by R. A. Neher and T. Leitner (28) (https://github.com/jnarag/HCVPacbioAnalysis/recombination.py). In brief, this method uses pairs of biallelic sites to examine how the frequency of recombinant haplotypes changes over time and distance between the sites.

## RESULTS

### Within-host population dynamics during HCV infection

Figure 1 summarises the within-host evolutionary dynamics of the four individuals from acute to chronic HCV infection. We observed a general tendency for HCV population genetic diversity to initially increase and subsequently stabilise in all individuals (Figure 1A). The within-host time-scaled phylogenies in three individuals (Figure 1B; p4, p37, and p61) were characterised by multiple viral lineages emerging in the first few years of infection, which were subsequently intermittently detected over the course of infection. This pattern was less strong in individual p37, and absent in individual p53, suggesting either not all viral lineages were sampled in the blood plasma, or the degree of within-host HCV population diversification varies among individuals. Viral sequences sampled from the same time point typically clustered in one or more clades of closely related sequences, giving rise to a burst of viral variants appearing in the phylogenies (Figure 1B). Together, these findings indicate infection in these individuals was likely initiated by a single viral strain, followed by viral diversification and establishment, leading to the emergence and maintenance of multiple co-circulating within-host HCV lineages.

**Figure 1:**
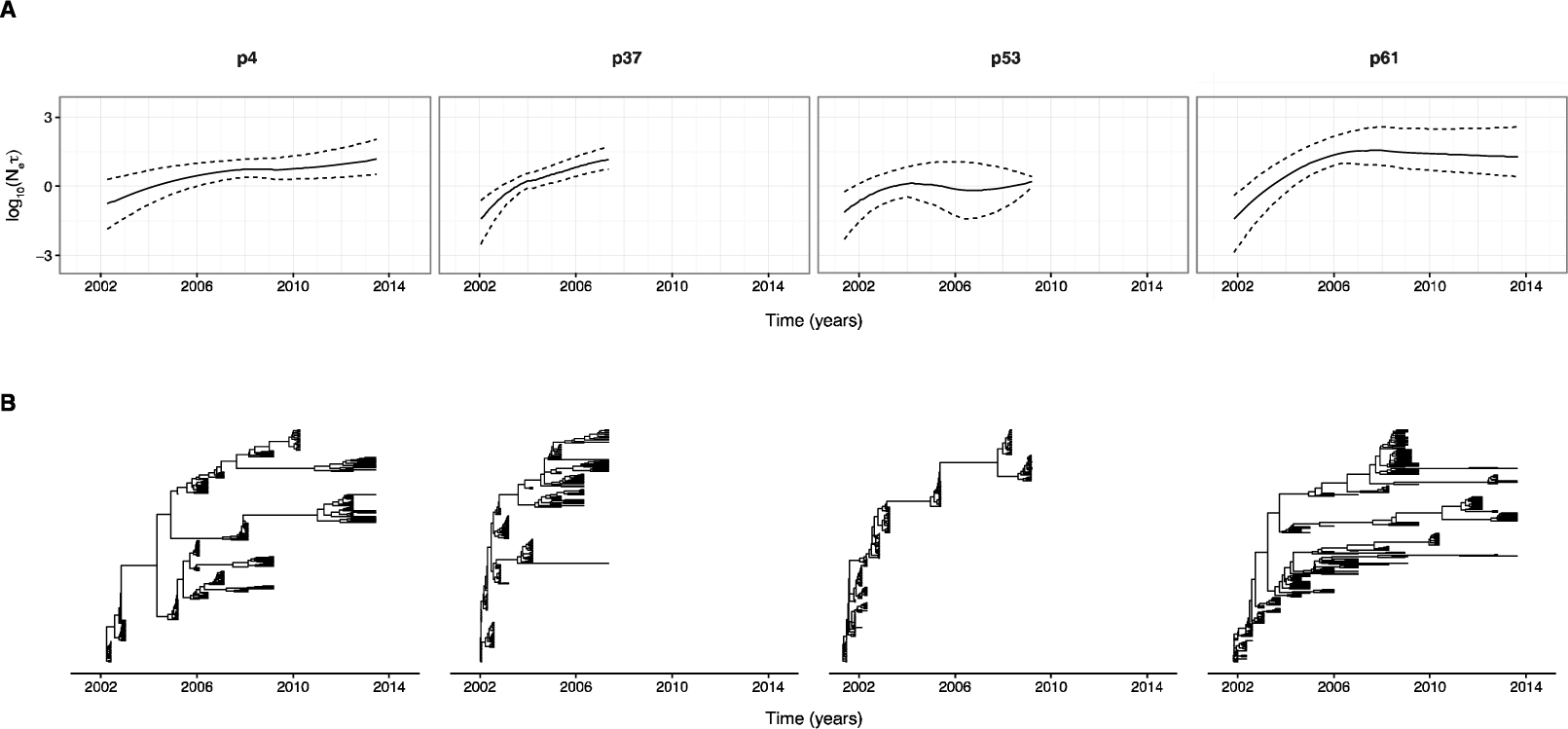
A) Within-host HCV population demographic histories and B) time-scaled phylogenies for the four HCV/HIV coinfected individuals. In A) the dashed lines correspond to loess regression fit through the lower and upper 95% highest posterior density estimates of N_e_τ, while the solid lines correspond to the smooth curve fitting to the mean N_e_τ over time. N_e_ corresponds to effective population size, while τ corresponds to the viral generation time, in calendar time.

Next, we examined whether patterns of HCV population genetic diversity were more consistent with a structured within-host population, or with a single population whose size varied through time. Support for these alternative hypotheses was evaluated by estimating their corresponding marginal likelihoods, which can be interpreted as the probability of observing the sequence data for a given population dynamic model. Consequently, if a comparatively higher marginal likelihood is found for one model over the other, this suggests there is stronger support from the data for that particular model. For three individuals (p37, p53, and p61), by comparing the marginal likelihoods between BASTA and the Bayesian skyline coalescent model, we found stronger evidence for a structured within-host HCV population, than for a single population with time-varying dynamics (Table 1). It is notable that despite the presence of multiple viral lineages co-existing for extended periods (∼9 years) in individual p4, we did not find significant support for a structured population. Interestingly, for individual p37, in spite of only one lineage being detected at any given sampling time point, our model comparison found greater evidence for a structured viral population (Table 1). One explanation for the lack of strong statistical support for a structured population in individual p4 could be that the structured coalescent model described by BASTA assumes a large, constant population size over time in each deme. This might not sufficiently capture the within-host HCV population dynamics in this individual, for example, if the viral subpopulations change in size over time, or if one is very small. Regardless, to determine the extent to which viral population structure affects HCV evolution during chronic infection, additional viral sequence data is required from a greater number of infected and untreated individuals.

**Table 1:** Marginal likelihoods estimated for BASTA and Bayesian Skyline using path-sampling

| **Individuals** | **BASTA** | **Bayesian skyline** |
| --- | --- | --- |
| p4 | -10279.3 | -10243.3 |
| p37 | -9508.2 | -9521.5 |
| p53 | -5900.1 | -5949.7 |
| p61 | -10277.3 | -10305.4 |

### Molecular evolution of the HCV envelope region

The codon partitioned substitution rates (CP1+2: 1st and 2nd codon positions; CP3: third codon position) for the three different gene regions within the envelope glycoprotein, E1, E2 excluding HVR1, and HVR1, are summarised in Table 2. The mean CP1+2 rates vary considerably both among gene regions and among individuals, with the HVR1 exhibiting the highest rate of evolution, which ranged from 1.5 to 4.8 x 10^-2^ subs/site/year. In individual p4, an elevated evolutionary rate is observed in the E1 region, corresponding to an accumulation of substitutions at 1^st^ and 2^nd^ codon positions that is 7 to 10 times faster than observed in the other individuals (Table 2). A similar difference in evolutionary rates is also observed in the E2 region, with individuals p4 and p37 accumulating substitutions 5 to 11 times faster than p53 and p61. As noted previously, the fastest evolving region within the HCV envelope region is the HVR1 (8, 14, 29, 30), which is consistent with stronger diversifying selection acting upon this region, most likely reflecting its role as a target of humoral immunity.

**Table 2:** Mean within-host evolutionary rates in HCV envelope glycoprotein. 95% credible intervals are given in brackets.

| Gene Regions | Partition | Within-host evolutionary rates ($10^{-2}$ subs/site/year) | | | |
| --- | --- | --- | --- | --- | --- |
|  |  | Individuals |  |  |  |
|  |  | p4 | p37 | p53 | p61 |
| E1 | CP1+2 | 1.942<br>(1.467, 2.465) | 0.193<br>(0.127, 0.257) | 0.240<br>(0.132, 0.347) | 0.211<br>(0.146, 0.278) |
| E2 | CP1+2 | 1.791<br>(1.303, 2.248) | 2.233<br>(1.602, 2.834) | 0.363<br>(0.237, 0.504) | 0.209<br>(0.159, 0.265) |
| HVR1 | CP1+2 | 3.361<br>(1.805, 6.255) | 4.841<br>(2.855, 6.727) | 3.540<br>(1.113, 8.772) | 1.475<br>(1.115, 1.921) |
| E1 | CP3 | 0.476<br>(0.224, 0.747) | 0.187<br>(0.119, 0.255) | 0.202<br>(0.109, 0.295) | 0.168<br>(0.129, 0.210) |
| E2 | CP3 | 0.572<br>(0.425, 0.722) | 0.443<br>(0.317, 0.584) | 0.221<br>(0.135, 0.314) | 0.257<br>(0.210, 0.313) |
| HVR1 | CP3 | 0.726<br>(0.272, 1.123) | 2.584<br>(0.927, 8.089) | 0.896<br>(0.191, 1.865) | 0.592<br>(0.271, 0.964) |
| Envelope | Total | 2.390<br>(2.014, 2.824) | 2.514<br>(2.030, 3.027) | 0.749<br>(0.503, 1.037) | 0.746<br>(0.650, 0.841) |

Differences in CP3 rates are in general less marked, with the exception of a higher substitution rate in the HVR1 observed for individual p37 (although this is associated with large statistical uncertainty; Table 2). The overall substitution rates of the envelope gene regions are also shown in Table 2; these suggest that the within-host HCV populations in the four individuals have evolved at varying rates, with slower rates observed in individuals p53 and p61 than in individuals p4 and p37. These differences probably result from individual variation in host immune responses and the underlying ecology of the within-host viral population (e.g. the number of viral subpopulations established during infection).

Figure 2 illustrates the per-site amino-acid diversity in the envelope glycoprotein in the four individuals. Most of the E1 and E2 gene regions are conserved in all four individuals. As expected, the HVR1 is characterised by the greatest amino-acid diversity, although there are regions in E2, and to a lesser extent in E1, which have comparable levels of per-site amino-acid diversity. There is some evidence of elevated amino-acid diversity at the nAb epitope located at position 249-269 aa in the E2 region (H77 reference position 428-447aa) in the four individuals, where the per-site amino-acid diversity is ∼0.25 or greater in 3 out of 4 individuals. However, without information on the specific immune response elicited in these individuals, it is difficult to determine if this pattern is driven by viral immune evasion. Interestingly, in individual p37, two regions outside of HVR1 in the E2 glycoprotein exceed a per-site amino-acid diversity of 0.25 (Figure 2; amino-acid positions 288 and 398, respectively), which might explain the comparatively higher CP1+2 rate observed in the E2 region of this individual (Figure 2 and Table 2). Interestingly, site 398 falls within an N-linked glycosylation site, which exhibits considerable variation during infection, possibly reflecting ongoing viral evasion from host immune responses.

**Figure 2.**
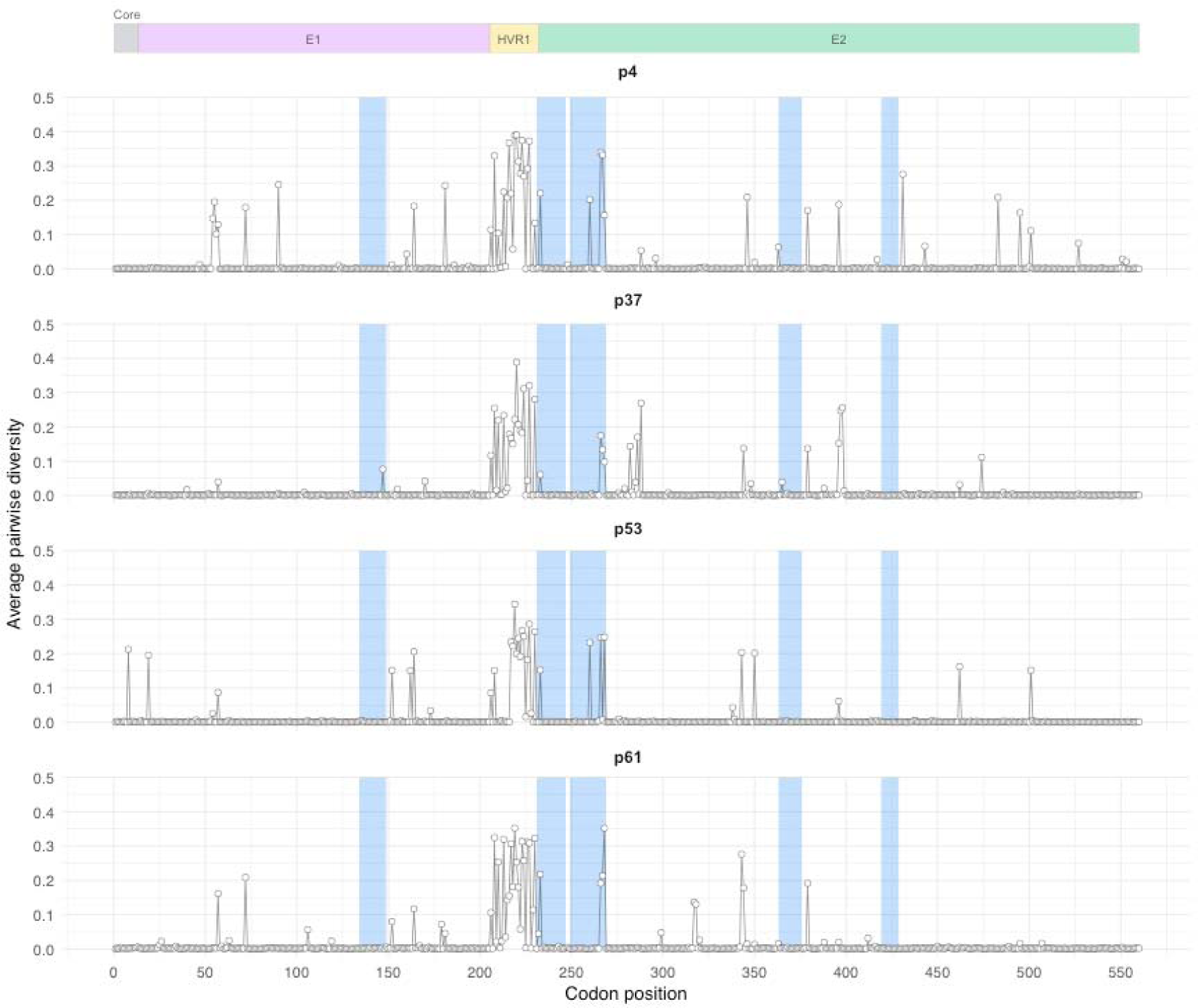
Mean amino-acid diversity per site. The genomic regions (Core, E1, HVR1, and E2) are indicated above the plot. In addition, the shaded blue regions indicate putative neutralizing antibody (nAb) epitopes. This information was collated from the Immune Epitope Database, which corresponds to regions in HCV (irrespective of genotype or subtype) that have been experimentally confirmed to elicit an antibody response.

### Population genetic analysis of HCV infection

We also analysed the full sequence dataset obtained for each individual (ranging from 5479 to 9122 sequences across all time points per individual) using simpler, population genetic summary statistics. First, we measured the divergence of the within-host viral population in three different gene regions for all individuals (Figure 3). The overall patterns of divergence are consistent with the rates of evolution estimated in Table 2. In general, the E1 and E2 regions accumulated nonsynonymous and synonymous substitutions at similar rates within individuals; while in HVR1 the nonsynonymous divergence was an order of magnitude higher. This supports the hypothesis that the HVR1 is mainly experiencing immune-mediated selection and the E1 and E2 gene regions are mostly subjected to purifying selection (Figure 3). Furthermore, while the gene regions largely diverged at an approximately constant rate over time, in individuals p4 and p61 we found that divergence of the HVR1 slowed down after 1 to 2 years of infection (Figure 3).

**Figure 3:**
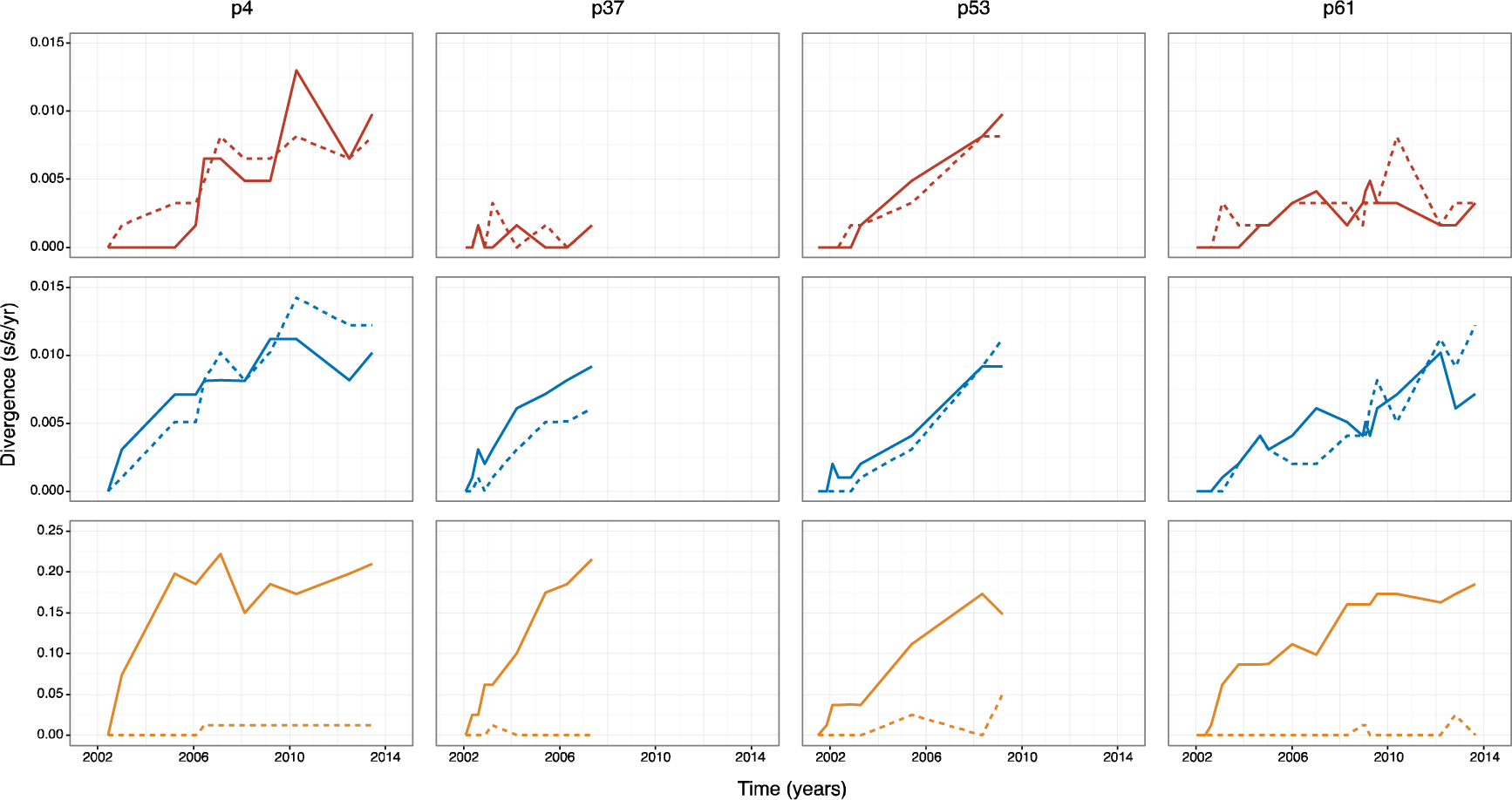
Divergence over time for each gene region (E1, E2, and HVR1 in red, blue, and yellow, respectively). Solid lines correspond to nonsynonymous divergence, while dashed lines correspond to synonymous divergence

The average within-host recombination rate during HCV infection, across all individuals was estimated to be 0.28 x 10^-7^ recombination events per site per day (IQR: 0.13-1.05 x 10^-7^). This is significantly lower than the inferred within-host substitution rate, which ranged from 2.05 to 8.21 x 10^-5^ substitutions per site per day. Furthermore, in contrast to the within-host recombination rate for HIV-1, which was estimated using exactly the same method (28), the HCV recombination rate is approximately two orders of magnitude lower. The low rates of within-host HCV recombination estimated here indicate that viral evolutionary dynamics during infection are likely to be influenced by strong linkage effects, and is further consistent with the low frequency of HCV circulating recombinant forms observed at the epidemiological scale (31).

## DISCUSSION

By examining the evolution of full-length HCV envelope glycoprotein sequences from acute to chronic infection, we find that HCV chronic infection is best explained by independently replicating viral subpopulations (9, 11, 32) that are established from a single infecting viral strain, but not all of which are detected in the plasma at all time points. We also found marked variation in the rates of evolution across the different regions of the envelope, and among individuals, for both synonymous and nonsynonymous mutations, combined with a very low rate of within-host recombination.

Given that longitudinal sampling of both the liver and blood from untreated HCV-infected individuals is unlikely to be feasible, or ethical, it has been difficult to investigate directly if chronic HCV infection is maintained by a structured viral population. To address this using an evolutionary approach, we assessed formally whether the observed within-host HCV population dynamics in the four individuals support a structured viral population characterised by two demes, where only of one these subpopulations is observed in the blood at any point in time, or with a single population whose population size varies over time. This analysis was possible due to the comparatively long-reads in this dataset, which provided sufficient statistical power to evaluate the two alternative hypotheses of the underlying population dynamics of HCV infection. For three individuals, we found a statistically better fit for a structured viral population. Although this result overall supports our original hypothesis, it is unclear why we did not find evidence for a structured population in one of the individuals despite observing co-circulation of multiple viral lineages that are intermittently detected during chronic infection. The lack of evidence for a structured viral population could reflect more complex within-host HCV population dynamics than assumed by the structured coalescent, such as one or more viral subpopulations are undergoing changes in population size, or if only one viral subpopulation is predominantly observed during chronic HCV infection. Furthermore, if more than one viral subpopulation is detected in the blood, this may resemble more closely a single well-mixed population rather than a heterogeneous, structured population.

Molecular evolutionary analyses of the HCV envelope region revealed very high rates of nonsynonymous substitution at the HVR1 region, which is consistent with this small genomic region undergoing strong immune-mediated selection (14, 29, 30, 33, 34). In contrast, the E1 and E2 (excluding HVR1) regions are largely characterised by purifying selection, indicating strong functional constraints acting upon these gene regions. Although similar conclusions have been reported previously (8, 14, 29, 30), the combination of frequent sampling during HCV infection and long-read sequence data has enabled us to robustly compare the rates of molecular evolution among the different gene regions in the envelope glycoprotein both within and between individuals.

In this study we also report, for the first time, an estimate of the within-host recombination rate of HCV during infection, which can be directly compared with other evolutionary estimates. Specifically, we found the within-host HCV recombination rate to be four orders of magnitude lower than the overall substitution rate of the HCV envelope glycoprotein, indicating strong linkage effects shape viral evolutionary dynamics during infection. In comparison to HIV-1 (28), recombination appears to be a weak evolutionary force during HCV within-host evolution, which helps explain why considerably fewer circulating recombinant forms are observed at the epidemiological scale for HCV than for HIV-1, even though HCV is more transmissible and mixed infections with distinct genotypes are relatively common. Another implication of limited recombination is that strong linkage effects are likely to shape HCV within-host viral evolution. In particular, selection is expected to be less effective in non-recombining populations due to clonal interference (35) and background selection (36-38), consequently reducing the overall rate of fixation of beneficial mutations in the population. A structured population can also limit viral adaptation if migration rates between viral subpopulations are low (39), since beneficial mutations are likely to be restricted to the subpopulations in which they emerged, thus preventing them from sweeping through the global population. This suggests that while drug resistance mutations may emerge during infection, their likely fixation within individuals and transmission between individuals may be limited.

To fully understand the extent to which within-host HCV populations are structured, and the effect that this, combined with low rates of recombination, has on HCV evolution during infection, will require additional viral sequence data that has been serially sampled from a greater number of individuals. As well as collecting key clinical information, such as HLA background, viral load and antibody responses, greater priority should be given to long-read, deep-sequence data that spans the whole virus genome, since this will give the power needed to determine whether, and how many, distinct viral subpopulations exit. This will be especially important for determining if viral population structure is associated with strong selection or genetic drift, and will help to elucidate the relative contribution of cell-mediated and humoral immunity during HCV infection.

